# Anti- SARS-CoV-2 Receptor Binding Domain Antibody Evolution after mRNA Vaccination

**DOI:** 10.1101/2021.07.29.454333

**Authors:** Alice Cho, Frauke Muecksch, Dennis Schaefer-Babajew, Zijun Wang, Shlomo Finkin, Christian Gaebler, Victor Ramos, Melissa Cipolla, Pilar Mendoza, Marianna Agudelo, Eva Bednarski, Justin DaSilva, Irina Shimeliovich, Juan Dizon, Mridushi Daga, Katrina Millard, Martina Turroja, Fabian Schmidt, Fengwen Zhang, Tarek Ben Tanfous, Mila Jankovic, Thiago Y. Oliveria, Anna Gazumyan, Marina Caskey, Paul D. Bieniasz, Theodora Hatziioannou, Michel C. Nussenzweig

## Abstract

Severe acute respiratory syndrome coronavirus 2 (SARS-CoV-2) infection produces B-cell responses that continue to evolve for at least one year. During that time, memory B cells express increasingly broad and potent antibodies that are resistant to mutations found in variants of concern^1^. As a result, vaccination of coronavirus disease 2019 (COVID-19) convalescent individuals with currently available mRNA vaccines produces high levels of plasma neutralizing activity against all variants tested^1, 2^. Here, we examine memory B cell evolution 5 months after vaccination with either Moderna (mRNA-1273) or Pfizer-BioNTech (BNT162b2) mRNA vaccines in a cohort of SARS-CoV-2 naïve individuals. Between prime and boost, memory B cells produce antibodies that evolve increased neutralizing activity, but there is no further increase in potency or breadth thereafter. Instead, memory B cells that emerge 5 months after vaccination of naïve individuals express antibodies that are similar to those that dominate the initial response. While individual memory antibodies selected over time by natural infection have greater potency and breadth than antibodies elicited by vaccination, the overall neutralizing potency of plasma is greater following vaccination. These results suggest that boosting vaccinated individuals with currently available mRNA vaccines will increase plasma neutralizing activity but may not produce antibodies with breadth equivalent to those obtained by vaccinating convalescent individuals.

---

Between January 21 and July 20, 2021, we recruited 32 volunteers with no history of prior SARS-CoV-2 infection receiving either Moderna (mRNA-1273; n=8) or Pfizer-BioNTech (BNT162b2; n=24) mRNA vaccines for sequential blood donation. Matched samples were obtained at 2 or 3 time points. Individuals indicated as “prime” were sampled an average of 2.5 weeks after receiving their first vaccine dose. Individuals who completed their vaccination regimen were sampled after an average of 1.3 months after the boost (median=35.5 days) which is not statistically different from the 1.3 month sampling in our naturally infected cohort^3^ (median=38.5 days, p=0.21). Individuals sampled at 1.3 months were sampled again approximately 5 months after the second vaccine dose. The volunteers ranged in age from 23-78 years (median=34.5 years), 53% were male and 47% female (for details see Methods and Supplementary Tables 1 and 2).

## Plasma binding and neutralization assays

Plasma IgM, IgG, and IgA responses to SARS-CoV-2 receptor binding domain (RBD) were measured by enzyme linked immunosorbent assay (ELISA)^3^. As reported by others^2, 4–6^ there was a significant increase in IgG reactivity to RBD between prime and boost (p<0.0001, Fig. 1a). IgM and IgA titers were lower than IgG titers and remained low after the second vaccine dose (Extended data Fig. 1a and b). The magnitude of the response was inversely correlated with age after the prime (r=-0.54, p=0.005), but in this limited sample set the age difference was no longer significant at 1.3 or 5 months after the second vaccine dose (Fig. 1b, Extended data Fig. 1c). Between 1.3 and 5 months after the boost, anti-RBD titers of IgG and IgA decreased significantly. IgG titers decreased by an average of 4.3-fold (range: 1.7- to 10.2-fold) and the loss of activity was directly correlated to the time after vaccination (p<0.0001, Fig. 1a and c and Extended data Fig. 1a and b).

**Fig. 1:**
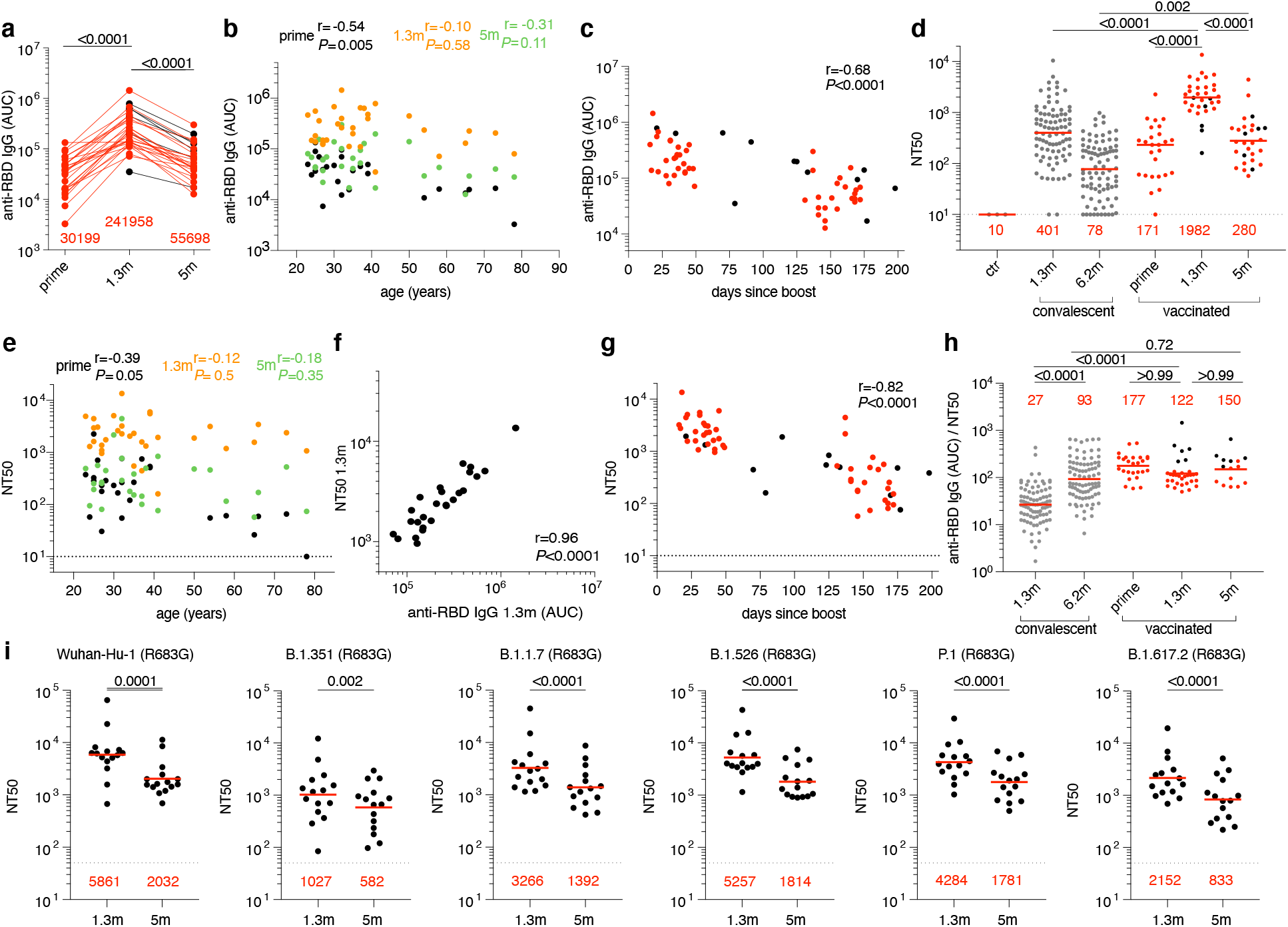
Plasma ELISAs and neutralizing activity. **a,** Graph shows area under the curve (AUC, Y-axis) for plasma IgG antibody binding to SARS- CoV-2 RBD after prime, and 1.3 months (m) and 5 months (m) post-second vaccination for n=32 paired samples. Samples without a prime value are shown in black (n=6). **b**, Graph shows plasma IgG antibody binding (AUC, Y-axis) plotted against age (X-axis) after prime (black), and 1.3 months (orange) and 5 months (green) post-second vaccination. **c**, Graph shows AUC values from **a** (Y-axis) plotted against time after vaccination (X-axis). Samples without a prime value are shown in black. **d,** NT50 values in pre-pandemic plasma (ctr) as well as plasma from convalescent individuals 1.3m^3^ and 6.2m^7^ after infection (grey) and in n=32 vaccinated individuals after prime, and 1.3- and 5-months after receiving 2 doses of an mRNA vaccine. Samples without a prime value are shown in black. **e**, NT50 values (Y-axis) vs. age (years, X-axis) in individuals after prime (black), or 1.3 months (1.3m, orange) or 5 months (5m, green) after boosting with an mRNA vaccine. **f**, NT50 values (Y-axis) vs. IgG antibody binding (AUC, X-axis) 1.3 months after 2 doses of an mRNA vaccine (n=26). **g,** Graph shows NT50 values (Y-axis) vs. days after boost (X-axis) in individuals receiving two doses of an mRNA vaccine. Samples without a prime value are shown in black. **h,** Ratio of anti-RBD IgG antibody (AUC) to NT50 values (Y-axis) plotted for convalescent infected individuals (grey) 1.3m^3^ or 6.2m^7^ after infection, and vaccinated individuals after the prime, and 1.3m and 5m after receiving 2 doses of an mRNA vaccine. Samples without a prime value are shown in black. **i**, Plasma neutralizing activity against indicated SARS-CoV-2 variants of interest/concern (n=15 paired samples at 1.3- and 5-months after full vaccination). Refer to Methods for a list of all substitutions/deletions/insertions in the spike variants. All experiments were performed at least in duplicate. Red bars and values in **a, d, h** and **i** represent geometric mean values. Statistical significance in **a, d, h** and **i** was determined by Kruskal-Wallis test with subsequent Dunn’s multiple comparisons, and in **b, c, e, f,** and **g** by Spearman correlation test.

Neutralizing activity was measured using HIV-1 pseudotyped with the SARS-CoV-2 spike^1, 3, 7, 8^. Naïve individuals showed variable responses to the initial vaccine dose with a geometric mean half-maximal neutralizing titer (NT50) of 171 (Fig. 1d, Supplementary Table 2). The magnitude of the neutralizing responses to the initial vaccine dose in naïve volunteers was inversely correlated with age (r=-0.39, p=0.05, Fig. 1e). Both binding and neutralizing responses to the second vaccine dose were correlated to the prime (r=0.46, p=0.02, Extended data. Fig. 1d; r=0.54, p=0.003, Extended data Fig. 1e) and produced a nearly 12-fold increase in the geometric mean neutralizing response that was similar in males and females and eliminated the age-related difference in neutralizing activity in the individuals in this cohort (Fig. 1d, Extended data Fig. 1f and Fig. 1e and Extended data Fig. 1g). 1.3 and 5 months after the boost naïve vaccinees had 4.9- and 3.6 fold higher neutralizing titers than a cohort of infected individuals measured 1.3^3^- and 6.2^7^-months after symptom onset, respectively (p<0.0001, Fig. 1d). Neutralizing responses were directly correlated to IgG anti-RBD titers (r=0.96, p<0.0001, Fig. 1f). Thus, the data obtained from this cohort agree with prior observations showing a significant increase in plasma neutralizing activity that are correlated with improved vaccine efficacy in naïve individuals that receive the second dose of mRNA vaccine^2, 6, 9, 10^ and higher neutralizing titers in fully vaccinated than infected individuals^2, 6^.

The 28 individuals assayed 5 months after vaccination had a 7.1-fold decrease in geometric mean neutralizing titer from their 1.3-month measurement (p<0.0001, Fig. 1d), with a range of 1.4- to 27-fold. Neutralizing activity was inversely correlated with the time from vaccination (r=-0.82, p<0.0001, Fig. 1g), and directly correlated to IgG anti-RBD binding titers when assessed 5 months after vaccination (Extended data. Fig. 1h). As reported by others^11^, the ratio of binding to neutralizing serum titers was significantly higher in vaccinated than convalescent individuals at the 1.3-month time point (p<0.0001, Fig. 1h). However, the difference was no longer apparent at the later time point (Fig. 1h).

We and others showed that the neutralizing responses elicited by mRNA vaccination are more potent against the original Wuhan Hu-1 strain than for some of the currently circulating variants of concern^2, 12–14^. To confirm these observations, we measured the neutralizing activity of 15 paired plasmas from naive individuals 1.3 and 5 months after the second vaccine dose against B.1.1.7 (alpha variant), B.1.351 (beta variant), B.1.526 (iota variant), P.1 (gamma variant) and B.1.617.2 (delta variant). Consistent with previous reports^13, 15–17^ the neutralizing activity against the variants was lower than against the original Wuhan Hu-1 strain (Fig. 1i, Supplementary Table 3). Initial geometric mean neutralizing titers at 1.3 months against B.1.351, B.1.1.7, B.1.526, P.1 and B.1.617.2 were 5.7, 1.8, 1.1, 1.4 and 2.7-fold lower than against Wuhan-Hu respectively (Fig. 1i). In the months following vaccination there was a decrease in neutralizing activity against Wuhan Hu-1 (R683G) and all the variants with geometric mean neutralizing titers for WT, B.1.351, B.1.1.7, B.1.526, P.1 and B.1.617.2 decreasing by 2.9-, 1.8-, 2.3-, 2.9-, 2.4- and 2.6-fold, respectively (Fig. 1i and Supplementary Table 3).

## Monoclonal Antibodies

Circulating antibodies produced by plasma cells can prevent infection if present at sufficiently high concentrations at the time of exposure. In contrast, the memory B cell compartment contains long lived antigen-specific B cells that mediate rapid recall responses that contribute to long term protection^18^. To examine the nature of the memory compartment elicited by one or two mRNA vaccine doses and its evolution after 5 months we used flow cytometry to enumerate B cells expressing receptors that bind to Wuhan Hu-1 (wild type, WT) and the B.1.351 K417N/E484K/N501Y variant RBDs (Fig. 2a and b, and Extended data Fig. 2). Although neutralizing antibodies develop to other parts of the spike (S) protein we focused on RBD because it is the dominant target of the memory antibody neutralizing response^19, 20^. Wuhan-Hu RBD-specific memory B cells developed after the prime in all volunteers examined and their numbers increased for up to 5 months after vaccination (Fig. 2a). Memory B cells binding to the B.1.351 K417N/E484K/N501Y variant RBD were detectable but in lower numbers than wild type RBD-binding B cells in all samples examined (Fig. 2b). Whereas IgG memory cells increased after the boost, IgM-expressing memory B cells that made up 23% of the memory compartment after the prime were nearly absent after boosting (Fig. 2c). Finally, circulating RBD-specific plasmablasts were readily detected after the prime but were infrequent after the boost (Fig. 2d, and Extended data Fig. 2d).

**Fig. 2:**
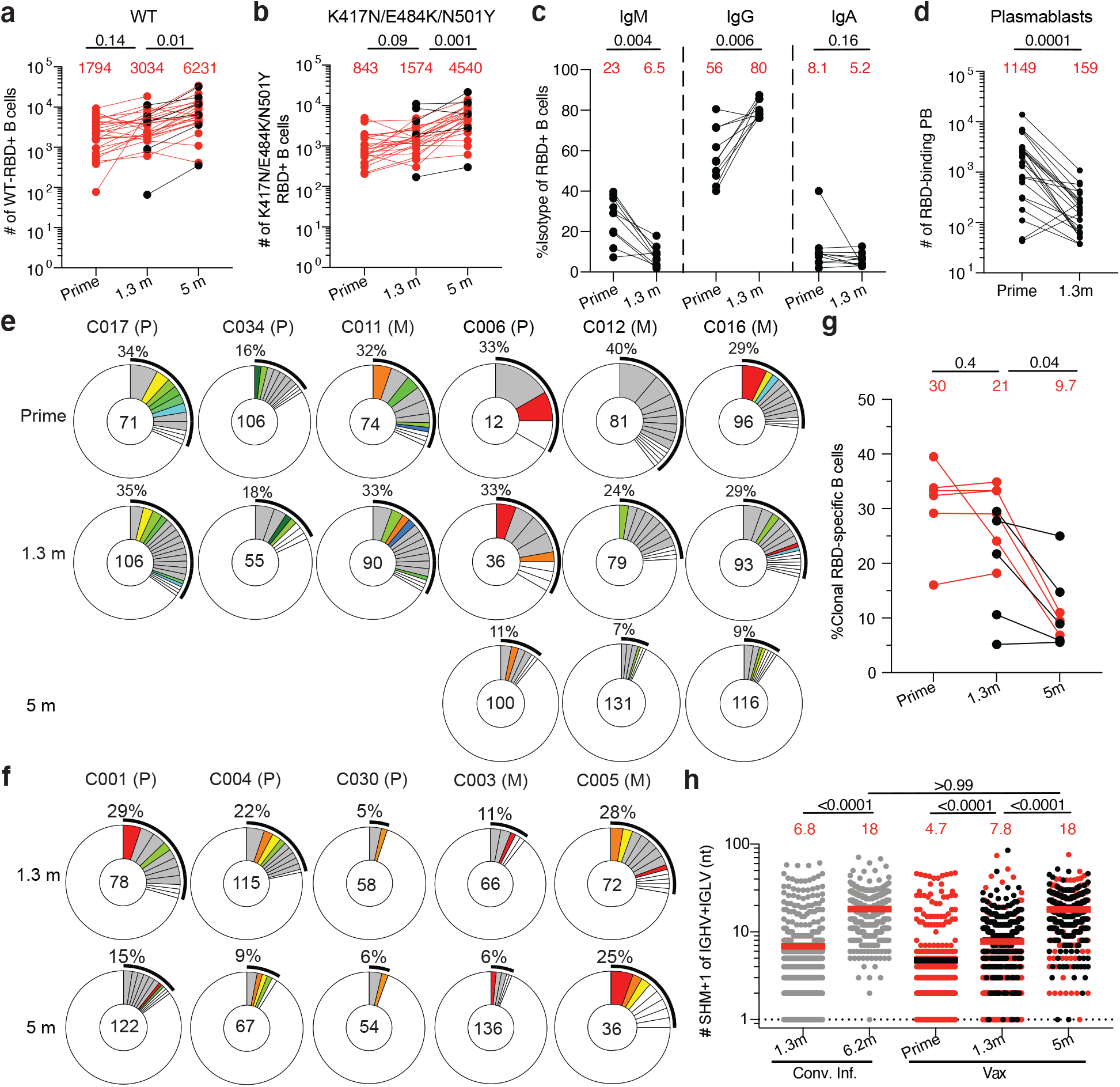
Anti-SARS-CoV-2 RBD B cells after vaccination. **a-d,** Graphs summarizing **a,** the number of Wuhan-Hu RBD (WT)-specific memory B cells per 10 million B cells for vaccinees and **b,** the number of antigen-specific memory B cells cross-reactive with both WT and K417N/E484K/N501Y RBD mutant per 10 million B cells for n=32 vaccinated individuals after prime, 1.3- and 5-months after 2 doses of vaccination. Samples without a prime value are shown in black. **c,** the frequency of IgM, IgG, or IgA isotype expression by Wuhan-Hu RBD-specific memory B cells after prime or 1.3 months post-boost (n=10), and **d,** number of Wuhan-Hu RBD-binding plasmablasts per 10 million B cells (n=26) after prime or 1.3 months post-boost. Red numbers indicate geometric means. Gating strategy is in Extended data Fig. 2. **e, and f,** Pie charts show the distribution of IgG antibody sequences obtained from memory B cells from 11 individuals after **e,** prime and 1.3-months or 5-months post-boost and **f,** 1.3- and 5-months post- boost. The number inside the circle indicates the number of sequences analyzed for the individual denoted above the circle, with Pfizer vaccinees indicated by (P) and Moderna by (M). Pie slice size is proportional to the number of clonally related sequences. The black outline and associated numbers indicate the percentage of clonally expanded sequences detected at each time point. Colored slices indicate persisting clones (same *IGHV* and *IGLV* genes, with highly similar CDR3s) found at more than one timepoint within the same individual. Grey slices indicate clones unique to the timepoint. White slices indicate repeating sequences isolated only once per time point. **g,** Graph shows the relative percentage of clonal sequences at each time point in **e** and **f**. The red numbers indicate the geometric means. Samples without a prime value are shown in black. **h,** Number of nucleotide somatic hypermutations (SHM) in the *IGHV* and *IGLV* combined (also Supplementary Table 4) in the antibodies illustrated in **e** and **f**, compared to the number of mutations obtained after 1.3^3^ or 6.2^7^ months after infection (grey). Horizontal bars and red numbers indicate mean number of nucleotide mutations at each time point. Samples without a prime value are shown in black. Statistical significance in **a, b, g,** and **h** was determined by Kruskal Wallis test with subsequent Dunn’s multiple comparisons, and in **c** and **d** was determined using Wilcoxon matched-pairs signed rank test.

The memory compartment continues to evolve up to one year after natural infection with selective enrichment of cells producing broad and potent neutralizing antibodies^1^. To determine how the memory compartment evolves after vaccination, we obtained 2328 paired antibody sequences from 11 individuals sampled at the time points described above (Fig. 2e and f, Extended data Fig 3, Supplementary Table 4). As expected *IGHV3-30* and *IGHV3-53* were over-represented after the first and second vaccine dose and remained over-represented 5 months after vaccination^21–23^ (Extended data Fig. 4).

All individuals examined showed expanded clones of memory B cells that expressed closely related *IGHV* and *IGHL* genes (Fig. 2e and f, Extended data Fig. 4). Paired prime and 1.3 month post boost samples showed expanded clones of memory B cells some of which were shared across plasmablasts, IgM and IgG prime, and IgG boost memory cells (Extended data Fig. 3 and 5). Thus, the cell fate decision controlling the germinal center versus plasmablast decision is not entirely affinity dependent since cells with the same initial affinity can enter both compartments to produce clonal relatives^24^.

The relative fraction of memory cells found in expanded clones varied between prime and boost and between individuals and decreased over time (Fig. 2e-g). Overall, clones represented 30%, 21%, and 9.7% of all sequences after prime, 1.3- and 5-month time points respectively (Fig. 2g). Nevertheless, clones of memory B cells continued to evolve for up to 5 months in vaccinated individuals as evidenced by the appearance of unique clones. Notably, unique clones appearing after 1.3 and 5 months represent a greater or equal fraction of the total memory B cell pool than the persisting clones (Fig. 2e-f, 16% vs 9.6% and 5.1% vs 4.7%, respectively, Extended data Fig. 3b). Finally, memory B cells emerging after the boost showed significantly higher levels of somatic mutations than plasmablasts or memory B cells isolated after the prime, and they continue to accumulate mutations up to 5 months post-boost (Fig. 2h, and Extended data Fig. 3d). In conclusion the memory B cell compartment continues to evolve for up to 5 months after mRNA vaccination.

## Neutralizing Activity of Monoclonal Antibodies

We performed ELISAs to confirm that the antibodies isolated from memory B cells bind to RBD (Extended data Fig. 6). 458 antibodies were tested by ELISA including: 88 isolated after the first vaccine dose; 210 isolated after the boost; and 160 isolated from individuals that had been fully vaccinated 5 months earlier. Among the 458 antibodies tested 430 (94%) bound to the Wuhan Hu-1 RBD indicating that the method used to isolate RBD-specific memory B cells was highly efficient (Supplementary Table 5-6). The geometric mean ELISA half-maximal concentration (EC50) of the antibodies obtained after prime, and 1.3- and 5-months after the second dose was 3.5, 2.9 and 2.7 ng/ml respectively, suggesting no major change in binding over time after vaccination (Extended data Fig. 6 and Supplementary Table 5-6).

430 RBD-binding antibodies were tested for neutralizing activity using HIV-1 pseudotyped with the SARS-CoV-2 spike^3, 8^. The geometric mean half-maximal inhibitory concentration (IC50) of the RBD-specific memory antibodies improved from 376 ng/ml to 153 ng/ml between the first and second vaccine dose (p=0.0005, Fig. 3a). The improvement was reflected in all clones (IC50 377 vs. 171 ng/ml, p=0.01 Fig. 3b), persisting clones (IC50 311 vs. 168, Fig. 3c, Supplementary Table 6), unique clones (IC50 418 vs. 165 ng/ml, p=0.03 Fig. 3d), and single antibodies (IC50 374 vs. 136 ng/ml, Fig. 3e). The increase in neutralizing activity between the first and second vaccine dose was associated with a decrease in the percentage of non-neutralizing antibodies (defined as IC50 >1000 ng/ml) and increased representation of neutralizing antibodies (p= 0.003, Fig. 3a). In conclusion, memory B cells recruited after the second dose account for most of the improvement in neutralizing activity in this compartment between the 2 vaccine doses. Thus, in addition to the quantitative improvement in serum neutralizing activity there is also an improvement in the neutralizing activity of the antibodies expressed in the memory compartment after boosting.

**Fig. 3:**
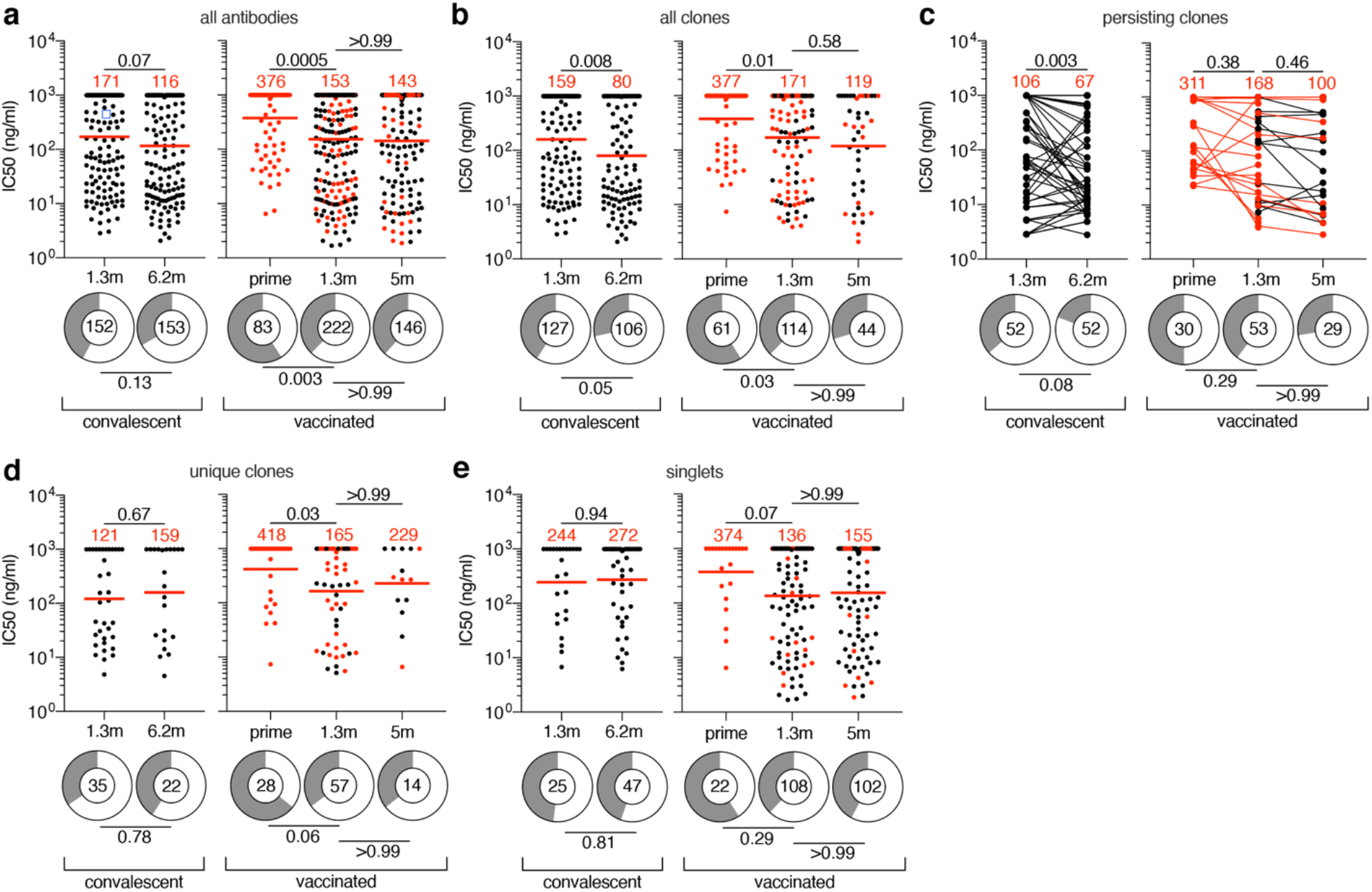
Anti-SARS-CoV-2 RBD monoclonal antibodies. **a-e,** Graphs show anti-SARS-CoV-2 neutralizing activity of monoclonal antibodies measured by a SARS-CoV-2 pseudotype virus neutralization assay using wild-type (Wuhan Hu-1^54^) SARS-CoV-2 pseudovirus^3, 8^. Half-maximal inhibitory concentration (IC50) values for all antibodies (**a**), all clones (**b**), persisting clones (**c**), unique clones (**d**) and singlets (**e**) isolated from COVID-19 convalescent individuals 1.3^3^ and 6.2^7^ months after infection or from vaccinated individuals after prime, and 1.3- or 5-months after 2 doses of vaccine. Each dot represents one antibody, where 451 total antibodies were tested including the 430 reported herein (Supplementary Table 5), and 21 previously reported antibodies derived from vaccinees within 8 weeks post-vaccination^13^. Antibodies isolated from samples without a prime value are shown in black. Pie charts illustrate the fraction of non-neutralizing (IC50 > 1000 ng/ml) antibodies (grey slices), inner circle shows the number of antibodies tested per group. Horizontal bars and red numbers indicate geometric mean values. Statistical significance was determined by Kruskal Wallis test with subsequent Dunn’s multiple comparisons. Statistical significance for ring plots was determined using Fisher’s exact test with subsequent Bonferroni-correction. All experiments were performed at least twice.

In contrast, there was no significant improvement in neutralizing activity of the monoclonal antibodies obtained between 1.3 and 5 months after vaccination (p>0.99, Fig. 3a). Although there was some improvement among B cell clones, which was accounted for by the small minority of persisting clones, neither was significant (p=0.58 and 0.46, Fig. 3b-e, Supplementary table 6). In contrast, memory antibodies obtained from convalescent individuals showed improved neutralizing activity between 1.3^3^ and 6.2 months^7^ with IC50 of 171 ng/ml to 116 ng/ml (Fig. 3a), which improved further after 1 year^1^. This improvement was due to increased neutralizing activity among persisting clones (p=0.003, Fig. 3c).

## Affinity, Epitopes and Neutralization Breadth

To examine affinity maturation after vaccination, we performed biolayer interferometry (BLI) experiments using the Wuhan Hu-1 RBD^3^. 147 antibodies were assayed, 30 obtained after the prime, 74 1.3-months after boosting, and 43 5-months after the boost. Geometric mean IC50s were comparable for the antibodies obtained from the 1.3- and 5-month time points (Extended data Fig. 7a). Overall, there was a 3- and 7.5 fold increase in affinity between the antibodies obtained between the first 2, and second 2 time points respectively (Fig. 4a). After 5 months the affinity of the antibodies obtained from vaccinated individuals was similar to antibodies obtained from naturally infected voluteers (Fig 4a). However, there was no correlation between affinity and neutralizing activity of the antibodies tested at any of the 3 time points (Extended data Fig. 7b).

**Fig. 4:**
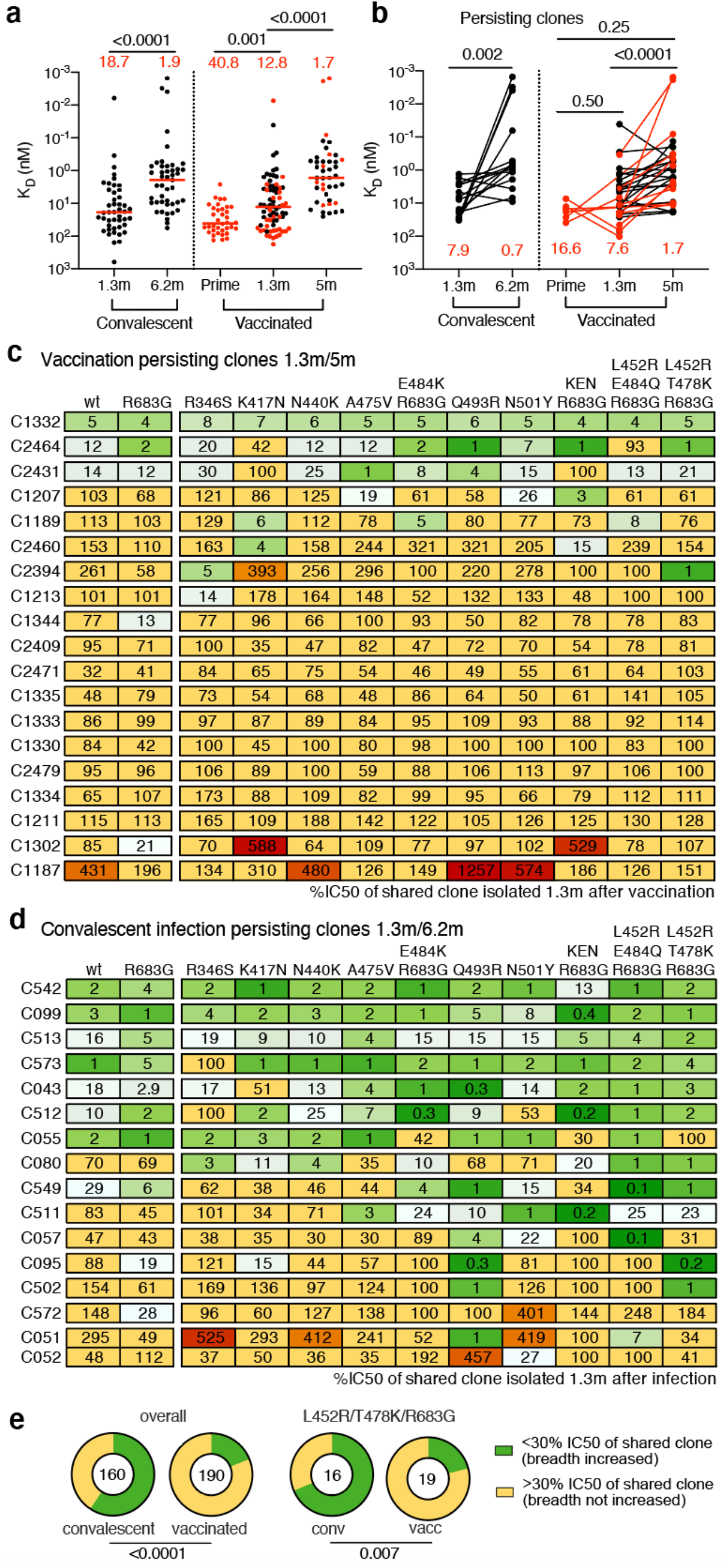
Affinity and Breadth. **a-b,** Graphs show antibody KDs for Wuhan-Hu RBD measured by BLI. **a,** antibodies isolated from convalescent individuals 1.3^3^- (n=42) and 6.2-months^7^ (n=45) after infection or from vaccinees after prime (n=36), and 1.3- (n=74) and 5-months (n=43) post- second vaccination. **b**, Clonally-paired antibodies isolated from convalescent individuals 1.3^3^- and 6.2^7^-months after infection (n=15) or vaccinated individuals between prime and 1.3 month (n=3), prime and 5 months (n=3), or 1.3- and 5-months after full vaccination (n=26). Antibodies isolated from samples without a prime value are shown in black. Red horizontal bars and numbers indicate median values. Statistical significance was determined using Kruskal Wallis test with subsequent Dunn’s multiple comparisons. **c-d,** Heat-maps show inhibitory concentrations of antibodies isolated 5m after vaccination (**c**) or 6.2m^7^ after infection (**d**) normalized to their shared clone isolated 1.3m after vaccination (**c**) or 1.3m^3^ after infection (**d**), expressed as %IC50, against indicated mutant SARS-CoV-2 pseudoviruses (Supplementary Table 8). Antibodies with improved (<30%) IC50 compared to their clonal relative isolated at an earlier timepoint are colored in shades of green with most improved antibodies in darkest green. Antibodies with worse (>300%) IC50 than their clonal relative isolated at an earlier timepoint are colored in red with the most worsened antibodies in dark red. Antibodies that did not change their IC50 by more than ∼3- fold are shown in yellow. **e,** Pie charts illustrate the fraction of antibodies showing improved (<30%, green) vs. not improved (yellow) IC50 compared to their clonal relative isolated at an earlier timepoint, inner circle shows the number of antibody-mutant combinations analyzed per group. Statistical significance for ring plots was determined using Fisher’s exact test.

We also compared the affinities of pairs of antibodies obtained from persisting clones between 1.3 and 5 months after vaccination. Persisting clones obtained at 1.3 and 5 months from vaccinated individuals showed a median 4.5-fold increase in affinity (p<0.0001, Fig. 4b). In contrast, a comparable group of persisting clonal antibodies obtained from convalescent individuals 1.3 and 6.2 months after infection showed a median 11.2-fold increase in affinity (p=0.002, Fig. 4b).

To determine whether the epitopes targeted by the monoclonal antibodies were changing over time, we performed BLI experiments in which a preformed antibody-RBD complex was exposed to a second monoclonal targeting one of 4 classes of structurally defined epitopes^1, 3^ (see schematic in Extended data Fig. 8a). There was no significant change in the distribution of targeted epitopes among 52 randomly selected antibodies with comparable neutralizing activity obtained from the 1.3- and 5-month time points (Extended data Fig. 8b and c, and Extended data Fig. 9).

In addition to the increase in potency, the neutralizing breath of memory antibodies obtained from persisting clones from convalescent individuals increases with time after infection^1, 7, 25^. To determine whether there is a similar increase in breadth with time after vaccination, we selected 20 random antibodies from the prime or 1.3 months after boost, with representative levels of activity against the original Wuhan Hu-1 strain, and measured their neutralization potency against a panel of pseudotypes encoding RBD mutations which were selected for resistance to different RBD antibody classes and/or are associated with circulating variants of concern (Extended data. Fig. 10). There was little change in breadth between prime and 1.3 months after boost, with only a small increase in resistance to K417N and A475V substitutions (Extended data Fig. 10, Supplementary Table 7).

In addition, we assayed 19 pairs of neutralizing antibodies expressed by persisting clones obtained 1.3 and 5 months after vaccination against the same mutant pseudotype viruses (Fig. 4c and Supplementary Table 8). They were compared to 7 previously reported^25^, plus 9 additional pairs of antibodies obtained from convalescent individuals at 1.3- and 6.2-month time points (Fig. 4d and Supplementary Table 8). Whereas only 36 of 190 (19%) of the vaccinee antibody- mutant combinations showed improved potency, 95 of the 160 (59%) convalescent pairs did so (p<0.0001, Fig. 4c, d and e). Moreover, only 4 of the 19 (21%) vaccine antibody pairs showed improved potency against pseudotypes carrying B.1.617.2 (delta variant)-specific RBD amino acid substitutions (L452R/T478K), while 11 out of 16 (69%) of the convalescent antibody pairs showed improved activity against this virus (p=0.007, Fig. 4c, d and e). We conclude that there is less increase in breadth in the months after mRNA vaccination than in a similar interval in naturally infected individuals.

Circulating antibodies are produced by an initial burst of short-lived plasmablasts^26–28^, and maintained by plasma cells with variable longevity^29–32^. SARS-CoV-2 infection or mRNA vaccination produces an early peak antibody response that decreases by 5-10-fold after 5 months^7, 33–37^. Notably, neutralization titres elicited by vaccination exceed those of COVID-19 recovered individuals at all comparable time points assayed. Nevertheless, neutralizing potency against variants is significantly lower than against Wuhan Hu-1, with up to 5-10-fold reduced activity against the B.1.351 variant^5, 6, 13, 14, 38^. Taken together with the overall decay in neutralizing activity there can be 1-2 orders of magnitude decrease in serum neutralizing activity after 5 or 6 months against variants when compared to the peak of neutralizing activity against Wuhan Hu-1. Thus, antibody mediated protection against variants is expected to wane significantly over a period of months, consistent with reports of reinfection in convalescent individuals and breakthrough infection by variants in fully vaccinated individuals^39–42^.

In contrast to circulating antibodies, memory B cells are responsible for rapid recall responses^43–,46^, and the number of cells in this compartment is relatively stable over the first 5-6 months after mRNA vaccination or natural infection^7, 47^. In both cases memory B cells continue to evolve as evidenced by increasing levels of somatic mutation and emergence of unique clones.

The memory response would be expected to protect individuals that suffer breakthrough infection from developing serious disease. Both natural infection and mRNA vaccination produce memory antibodies that evolve increased affinity. However, vaccine-elicited memory monoclonal antibodies show more modest neutralizing potency and breadth than those that developed after natural infection^1, 7^. Notably, the difference between the memory compartment that develops in response to natural infection vs mRNA vaccination reported above is consistent with the higher level of protection from variants conferred by natural infection^42^.

There are innumerable differences between natural infection and mRNA vaccination that could account for the differences in antibody evolution over time. These include but are not limited to: 1. Route of antigen delivery, respiratory tract vs. intra-muscular injection^48, 49^; 2. The physical nature of the antigen, intact virus vs. conformationally stabilized prefusion S protein^50^; 3. Antigen persistence, weeks in the case of natural infection^7^ vs. hours to days for mRNA^51^. Each of these could impact on B cell evolution and selection directly, and indirectly through differential T cell recruitment.

The increase in potency and breadth in the memory compartment that develops after natural infection accounts for the exceptional responses to Wuhan Hu-1 and its variants that convalescent individuals develop when boosted with mRNA vaccines^1, 5^. The expanded memory B cell compartment in mRNA vaccinees should also produce high titers of neutralizing antibodies when vaccinees are boosted or when they are re-exposed to the virus^52^. Boosting vaccinated individuals with currently available mRNA vaccines should produce strong responses that mirror or exceed their initial vaccine responses to Wuhan-Hu but with similarly decreased coverage against variants. Whether an additional boost with Wuhan-Hu-based or variant vaccines or re-infection will also elicit development of memory B cells expressing antibodies showing increased breadth remains to be determined. Finally, timing an additional boost for optimal responses depends on whether the objective is to prevent infection or disease^53^. Given the current rapid emergence of SARS-CoV-2 variants, boosting to prevent infection would likely be needed on a time scale of months. The optimal timing for boosting to prevent serious disease will depend on the stability and further evolution of the memory B cell compartment.

## METHODS

### Study participants

Participants were healthy volunteers receiving either the Moderna (mRNA-1273) or Pfizer-BioNTech (BNT162b2) mRNA vaccines against severe acute respiratory syndrome coronavirus 2 (SARS-CoV-2) who were recruited for serial blood donations at Rockefeller University Hospital in New York between January 21 and July 20, 2021. The majority of participants (n=28) were *de novo* recruited for this study, while a subgroup of individuals (n=4) were from a long-term study cohort^13^. Eligible participants were healthy adults with no history of infection with SARS-CoV-2, as determined by clinical history and confirmed through serology testing, receiving one of the two Moderna (mRNA-1273) or Pfizer-BioNTech (BNT162b2), according to current dosing and interval guidelines. Exclusion criteria included incomplete vaccination status, presence of clinical signs and symptoms suggestive of acute infection with or a positive reverse transcription polymerase chain reaction (RT-PCR) results for SARS-CoV-2 in saliva, or a positive (coronavirus disease 2019) COVID-19 serology. Seronegativity for COVID-19 was established through the absence of serological activity toward the nucleocapsid protein (N) of SARS-CoV-2. Participants presented to the Rockefeller University Hospital for blood sample collection and were asked to provide details of their vaccination regimen, possible side effects, comorbidities and possible COVID-19 history. All participants provided written informed consent before participation in the study and the study was conducted in accordance with Good Clinical Practice. The study was performed in compliance with all relevant ethical regulations and the protocol (DRO-1006) for studies with human participants was approved by the Institutional Review Board of the Rockefeller University. For detailed participant characteristics see Supplementary Tables 1 and 2.

### Blood samples processing and storage

Peripheral Blood Mononuclear Cells (PBMCs) obtained from samples collected at Rockefeller University were purified as previously reported by gradient centrifugation and stored in liquid nitrogen in the presence of Fetal Calf Serum (FCS) and Dimethylsulfoxide (DMSO)^3, 7^. Heparinized plasma and serum samples were aliquoted and stored at -20°C or less. Prior to experiments, aliquots of plasma samples were heat-inactivated (56°C for 1 hour) and then stored at 4°C.

### ELISAs

Enzyme-Linked Immunosorbent Assays (ELISAs)^55, 56^ to evaluate antibodies binding to SARS- CoV-2 RBD were performed by coating of high-binding 96-half-well plates (Corning 3690) with 50 μl per well of a 1μg/ml protein solution in Phosphate-buffered Saline (PBS) overnight at 4°C. Plates were washed 6 times with washing buffer (1× PBS with 0.05% Tween-20 (Sigma- Aldrich)) and incubated with 170 μl per well blocking buffer (1× PBS with 2% BSA and 0.05% Tween-20 (Sigma)) for 1 hour at room temperature. Immediately after blocking, monoclonal antibodies or plasma samples were added in PBS and incubated for 1 hour at room temperature. Plasma samples were assayed at a 1:66 starting dilution and 10 additional threefold serial dilutions. Monoclonal antibodies were tested at 10 μg/ml starting concentration and 10 additional fourfold serial dilutions. Plates were washed 6 times with washing buffer and then incubated with anti-human IgG, IgM or IgA secondary antibody conjugated to horseradish peroxidase (HRP) (Jackson Immuno Research 109-036-088 109-035-129 and Sigma A0295) in blocking buffer at a 1:5,000 dilution (IgM and IgG) or 1:3,000 dilution (IgA). Plates were developed by addition of the HRP substrate, 3,3’,5,5’-Tetramethylbenzidine (TMB) (ThermoFisher) for 10 minutes (plasma samples) or 4 minutes (monoclonal antibodies). The developing reaction was stopped by adding 50 μl of 1 M H2SO4 and absorbance was measured at 450 nm with an ELISA microplate reader (FluoStar Omega, BMG Labtech) with Omega and Omega MARS software for analysis. For plasma samples, a positive control (plasma from participant COV72, diluted 66.6- fold and ten additional threefold serial dilutions in PBS) was added to every assay plate for normalization. The average of its signal was used for normalization of all the other values on the same plate with Excel software before calculating the area under the curve using Prism V9.1(GraphPad). Negative controls of pre-pandemic plasma samples from healthy donors were used for validation (for more details please see^3^). For monoclonal antibodies, the ELISA half-maximal concentration (EC50) was determined using four-parameter nonlinear regression (GraphPad Prism V9.1). EC50s above 2000 ng/mL were considered non-binders.

### Proteins

The mammalian expression vector encoding the Receptor Binding-Domain (RBD) of SARS-CoV-2 (GenBank MN985325.1; Spike (S) protein residues 319-539) was previously described^57^.

### SARS-CoV-2 pseudotyped reporter virus

A panel of plasmids expressing RBD-mutant SARS-CoV-2 spike proteins in the context of pSARS-CoV-2-S Δ19 has been described^13, 25, 58^. Variant pseudoviruses resembling variants of interest/concern B.1.1.7 (first isolated in the UK), B.1.351 (first isolated in South-Africa), B.1.526 (first isolated in New York City), P.1 (first isolated in Brazil) and B.1.617.2 (first isolated in India) were generated by introduction of substitutions using synthetic gene fragments (IDT) or overlap extension PCR mediated mutagenesis and Gibson assembly. Specifically, the variant-specific deletions and substitutions introduced were:

B.1.1.7: ΔH69/V70, ΔY144, N501Y, A470D, D614G, P681H, T761I, S982A, D118H B.1.351: D80A, D215G, L242H, R246I, K417N, E484K, N501Y, D614G, A701V B.1.526: L5F, T95I, D253G, E484K, D614G, A701V.

P.1: L18F, T20N, P26S, D138Y, R190S, K417T, E484K, N501Y, D614G, H655Y, T1027I, V1167F B.1.617.2: T19R, Δ156-158, L452R, T478K, D614G, P681R, D950N

The E484K, K417N/E484K/N501Y, L452R/E484Q and L452R/T478K substitution, as well as the deletions/substitutions corresponding to variants of concern listed above were incorporated into a spike protein that also includes the R683G substitution, which disrupts the furin cleaveage site and increases particle infectivity. Neutralizing activity against mutant pseudoviruses were compared to a wildtype (WT) SARS-CoV-2 spike sequence (NC_045512), carrying R683G where appropriate.

SARS-CoV-2 pseudotyped particles were generated as previously described^3, 8^. Briefly, 293T cells were transfected with pNL4-3ΔEnv-nanoluc and pSARS-CoV-2-SΔ19, particles were harvested 48 hours post-transfection, filtered and stored at -80°C.

### Pseudotyped virus neutralization assay

Fourfold serially diluted pre-pandemic negative control plasma from healthy donors, plasma from COVID-19-convalescent individuals or monoclonal antibodies were incubated with SARS-CoV-2 pseudotyped virus for 1 hour at 37 °C. The mixture was subsequently incubated with 293TAce2 cells^3^ (for all WT neutralization assays) or HT1080Ace2 cl14 (for all mutant panels and variant neutralization assays) cells^13^ for 48 hours after which cells were washed with PBS and lysed with Luciferase Cell Culture Lysis 5× reagent (Promega). Nanoluc Luciferase activity in lysates was measured using the Nano-Glo Luciferase Assay System (Promega) with the Glomax Navigator (Promega). The relative luminescence units were normalized to those derived from cells infected with SARS-CoV-2 pseudotyped virus in the absence of plasma or monoclonal antibodies. The half-maximal neutralization titers for plasma (NT50) or half-maximal and 90% inhibitory concentrations for monoclonal antibodies (IC50 and IC90) were determined using four-parameter nonlinear regression (least squares regression method without weighting; constraints: top=1, bottom=0) (GraphPad Prism).

### Biotinylation of viral protein for use in flow cytometry

Purified and Avi-tagged SARS-CoV-2 RBD or SARS-CoV-2 RBD K417N/E484K/N501Y mutant was biotinylated using the Biotin-Protein Ligase-BIRA kit according to manufacturer’s instructions (Avidity) as described before^3^. Ovalbumin (Sigma, A5503-1G) was biotinylated using the EZ-Link Sulfo-NHS-LC-Biotinylation kit according to the manufacturer’s instructions (Thermo Scientific). Biotinylated ovalbumin was conjugated to streptavidin-BV711 (BD biosciences, 563262) and RBD to streptavidin-PE (BD Biosciences, 554061) and streptavidin- AF647 (Biolegend, 405237)^3^.

### Flow cytometry and single cell sorting

Single-cell sorting by flow cytometry was described previously^3^. Briefly, peripheral blood mononuclear cells were enriched for B cells by negative selection using a pan-B-cell isolation kit according to the manufacturer’s instructions (Miltenyi Biotec, 130-101-638). The enriched B cells were incubated in Flourescence-Activated Cell-sorting (FACS) buffer (1× PBS, 2% FCS, 1 mM ethylenediaminetetraacetic acid (EDTA)) with the following anti-human antibodies (all at 1:200 dilution): anti-CD20-PECy7 (BD Biosciences, 335793), anti-CD3-APC-eFluro 780 (Invitrogen, 47-0037-41), anti-CD8-APC-eFluor 780 (Invitrogen, 47-0086-42), anti-CD16-APC-eFluor 780 (Invitrogen, 47-0168-41), anti-CD14-APC-eFluor 780 (Invitrogen, 47-0149-42), as well as Zombie NIR (BioLegend, 423105) and fluorophore-labeled RBD and ovalbumin (Ova) for 30 min on ice. Single CD3-CD8-CD14-CD16−CD20+Ova−RBD-PE+RBD-AF647+ B cells were sorted into individual wells of 96-well plates containing 4 μl of lysis buffer (0.5× PBS, 10 mM Dithiothreitol (DTT), 3,000 units/ml RNasin Ribonuclease Inhibitors (Promega, N2615) per well using a FACS Aria III and FACSDiva software (Becton Dickinson) for acquisition and FlowJo for analysis. The sorted cells were frozen on dry ice, and then stored at −80 °C or immediately used for subsequent RNA reverse transcription. For plasmablast single-cell sorting, in addition to above antibodies, B cells were also stained with anti-CD19-BV605 (Biolegend, 302244), and single CD3-CD8-CD14-CD16-CD19+CD20-Ova-RBD-PE+RBD-AF647+ plasmablasts were sorted as described above. For B cell phenotype analysis, in addition to above antibodies, B cells were also stained with following anti-human antibodies: anti-IgD-BV421 (Biolegend, 348226), anti-CD27-FITC (BD biosciences, 555440), anti-CD19-BV605 (Biolegend, 302244), anti-CD71- PerCP-Cy5.5 (Biolegend, 334114), anti- IgG-PECF594 (BD biosciences, 562538), anti-IgM-AF700 (Biolegend, 314538), anti-IgA-Viogreen (Miltenyi Biotec, 130-113-481).

### Antibody sequencing, cloning and expression

Antibodies were identified and sequenced as described previously^3, 59^. In brief, RNA from single cells was reverse-transcribed (SuperScript III Reverse Transcriptase, Invitrogen, 18080-044) and the cDNA was stored at −20 °C or used for subsequent amplification of the variable IGH, IGL and IGK genes by nested PCR and Sanger sequencing. Sequence analysis was performed using MacVector. Amplicons from the first PCR reaction were used as templates for sequence- and ligation-independent cloning into antibody expression vectors. Recombinant monoclonal antibodies were produced and purified as previously described^3^.

### Biolayer interferometry

Biolayer interferometry assays were performed as previously described^3^. Briefly, we used the Octet Red instrument (ForteBio) at 30 °C with shaking at 1,000 r.p.m. Affinity measurement of anti-SARS-CoV-2 IgGs binding were corrected by subtracting the signal obtained from traces performed with IgGs in the absence of WT RBD. The kinetic analysis using protein A biosensor (ForteBio 18-5010) was performed as follows: (1) baseline: 60sec immersion in buffer. (2) loading: 200sec immersion in a solution with IgGs 10 μg/ml. (3) baseline: 200sec immersion in buffer. (4) Association: 300sec immersion in solution with WT RBD at 20, 10 or 5 μg/ml (5) dissociation: 600sec immersion in buffer. Curve fitting was performed using a fast 1:1 binding model and the Data analysis software (ForteBio). Mean equilibrium dissociation constant (KD) values were determined by averaging all binding curves that matched the theoretical fit with an R^2^ value ≥ 0.8.

### Computational analyses of antibody sequences

Antibody sequences were trimmed based on quality and annotated using Igblastn v.1.14. with IMGT domain delineation system. Annotation was performed systematically using Change-O toolkit v.0.4.540^60^. Heavy and light chains derived from the same cell were paired, and clonotypes were assigned based on their V and J genes using in-house R and Perl scripts. All scripts and the data used to process antibody sequences are publicly available on GitHub (https://github.com/stratust/igpipeline/tree/igpipeline2_timepoint_v2).

The frequency distributions of human V genes in anti-SARS-CoV-2 antibodies from this study was compared to 131,284,220 IgH and IgL sequences generated by^61^ and downloaded from cAb- Rep^62^, a database of human shared BCR clonotypes available at https://cab-rep.c2b2.columbia.edu/. Based on the 112 distinct V genes that make up the 7936 analyzed sequences from Ig repertoire of the 11 participants present in this study, we selected the IgH and IgL sequences from the database that are partially coded by the same V genes and counted them according to the constant region. The frequencies shown in Extended data Fig. 4 are relative to the source and isotype analyzed. We used the two-sided binomial test to check whether the number of sequences belonging to a specific *IGHV* or *IGLV* gene in the repertoire is different according to the frequency of the same IgV gene in the database. Adjusted p-values were calculated using the false discovery rate (FDR) correction. Significant differences are denoted with stars.

Nucleotide somatic hypermutation and Complementarity-Determining Region (CDR3) length were determined using in-house R and Perl scripts. For somatic hypermutations, *IGHV* and *IGLV* nucleotide sequences were aligned against their closest germlines using Igblastn and the number of differences were considered nucleotide mutations. The average number of mutations for V genes was calculated by dividing the sum of all nucleotide mutations across all participants by the number of sequences used for the analysis.

## Data availability statement

Data are provided in Supplementary Tables 1-8. The raw sequencing data and computer scripts associated with Figure 2 and Extended data Fig. 3 have been deposited at Github (https://github.com/stratust/igpipeline/tree/igpipeline2_timepoint_v2). This study also uses data from “A Public Database of Memory and Naive B-Cell Receptor Sequences” (https://doi.org/10.5061/dryad.35ks2), PDB (6VYB and 6NB6) and from “High frequency of shared clonotypes in human B cell receptor repertoires” (https://doi.org/10.1038/s41586-019-0934-8).

## Code availability statement

Computer code to process the antibody sequences is available at GitHub (https://github.com/stratust/igpipeline/tree/igpipeline2_timepoint_v2).

## Data presentation

Figures arranged in Adobe Illustrator 2020.

## Acknowledgments

We thank all study participants who devoted time to our research; The Rockefeller University Hospital nursing staff and Clinical Research Support Office and nursing staff. Mayu Okawa Frank, Marissa Bergh, and Robert B. Darnell for SARS-CoV-2 saliva PCR testing. Charles M. Rice, and all members of the M.C.N. laboratory for helpful discussions, Maša Jankovic for laboratory support, and Kristie Gordon for technical assistance with cell-sorting experiments. This work was supported by NIH grant P01-AI138398-S1 (M.C.N.) and 2U19AI111825 (M.C.N.). R37-AI64003 to P.D.B.; R01AI78788 to T.H. F.M. is supported by the Bulgari Women & Science Fellowship in COVID-19 Research. C.G. was supported by the Robert S. Wennett Post-Doctoral Fellowship. D.S.B and C.G. were supported in part by the National Center for Advancing Translational Sciences (National Institutes of Health Clinical and Translational Science Award program, grant UL1 TR001866), and C.G. by the Shapiro-Silverberg Fund for the Advancement of Translational Research. P.D.B. and M.C.N. are Howard Hughes Medical Institute Investigators.

## Author Contributions

P.D.B., T.H., and M.C.N. conceived, designed and analyzed the experiments. M. Caskey and C.G. designed clinical protocols. A.C, F.M., D.S.B., Z.W., S.F., P.M., M.A., E.B., J.D.S., I.S., J.D. F.S., F.Z., and T.B.T. carried out experiments. A.G. and M. Cipolla produced antibodies. D.S.B., M.D., M.T., K.G.M., C.G. and M. Caskey recruited participants, executed clinical protocols. T.Y.O. and V.R. performed bioinformatic analysis. A.C., F.M, D.S.B., Z.W., S.F., and M.C.N. wrote the manuscript with input from all co-authors.

## EXTENDED FIGURES

**Extended Data Fig 1:**
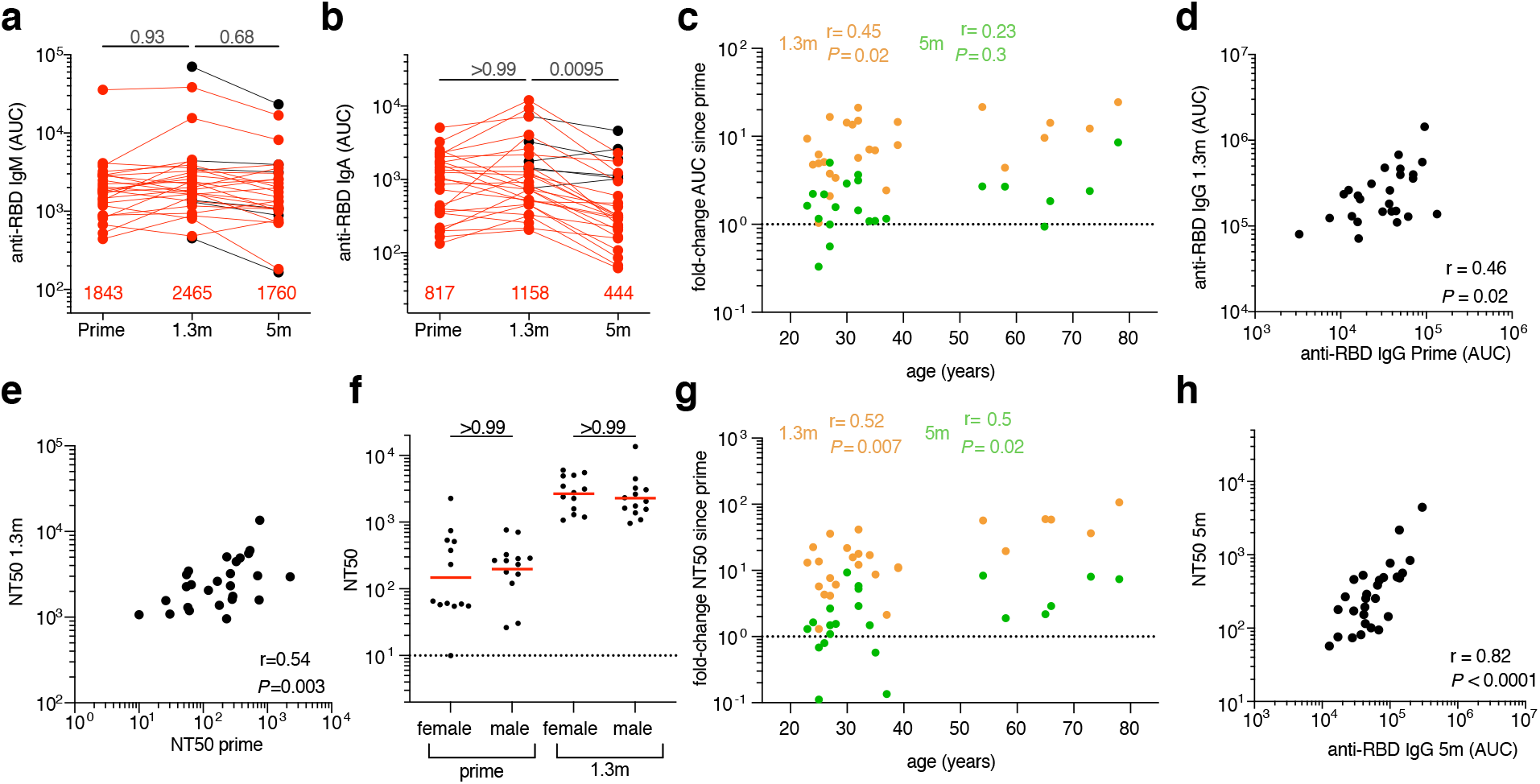
Plasma ELISA and neutralization. **a,b**, Graph shows area under the curve (AUC, Y-axis) for plasma IgM (**a**) or IgA (**b**) antibody binding to SARS-CoV-2 RBD after prime, and 1.3- and 5-months post-boost for paired samples from n=32 vaccinated individuals. Samples without a prime value are shown in black. **c,** Graph shows age (years, X-axis) vs. fold-change of IgG-binding titers (AUC, Y-Axis) between prime and 1.3m (orange) or 5m (green) post-boost. **d,** IgG antibody binding after prime (AUC, X-axis) vs. IgG antibody binding after 1.3 months post-boost (AUC, Y-axis) and **e**, NT50 values after prime (X-axis) vs. NT50 values after 1.3 months post-boost (Y-axis) in individuals receiving two doses of an mRNA vaccine (n=26). **f,** NT50 values after prime and 1.3 months post-boost in females and males receiving 2 doses of an mRNA vaccine. **g,** Graph shows age (years, X-axis) vs fold-change of NT50 (X-axis) between prime and 1.3m (orange) or 5m (green) post-boost. **h,** NT50 values (Y-axis) vs. IgG antibody binding (AUC, X-axis) 5 months after boost in individuals receiving two doses of an mRNA vaccine (n=28). All experiments were performed at least in duplicate. Red values or bar in **a**, **b** and **f** represent geometric mean values. Statistical significance in **a, b,** and **f** was determined by Kruskal-Wallis test with subsequent Dunn’s multiple comparisons, or by Spearman correlation test in **c, d, e, g,** and **h**.

**Extended Data Fig 2:**
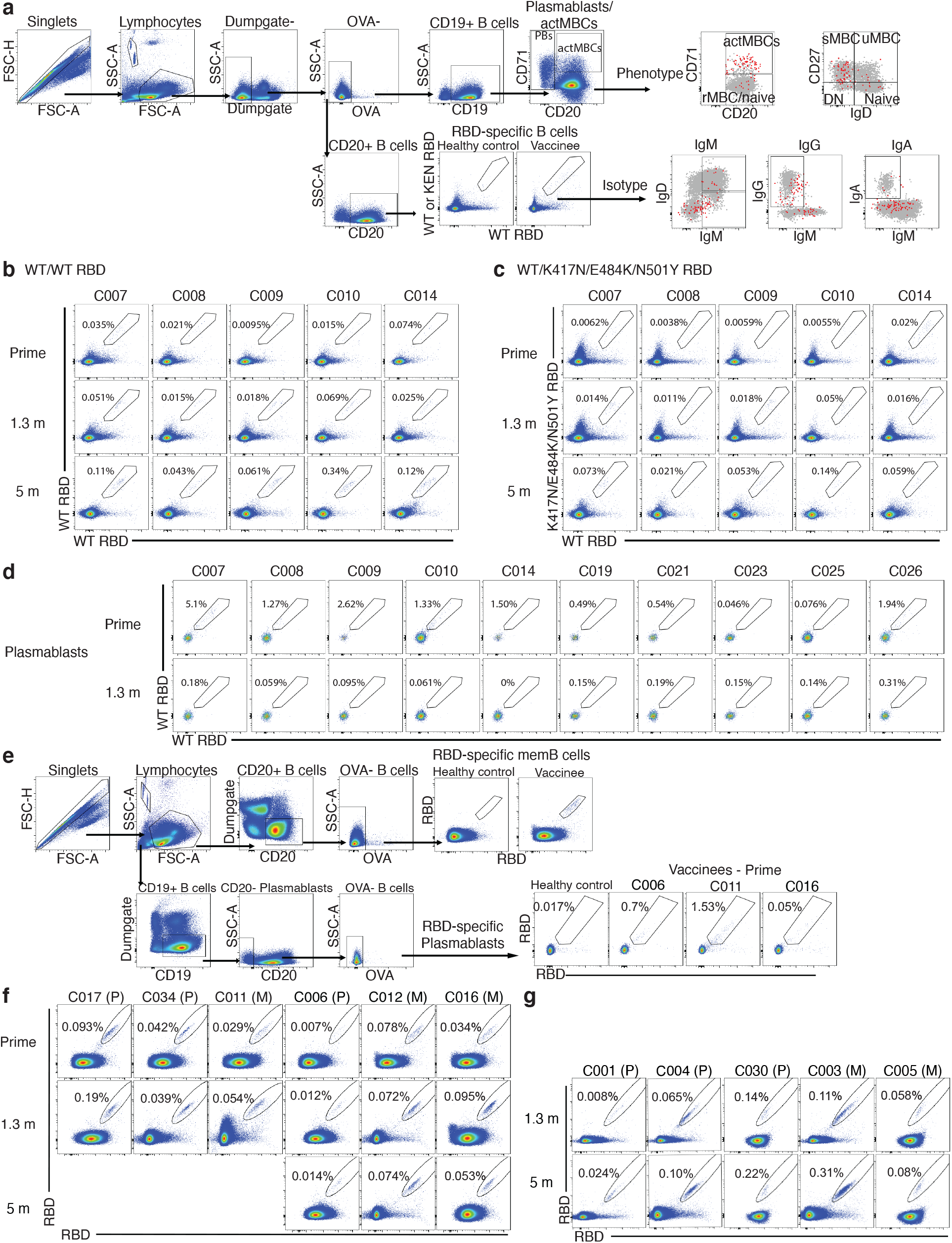
Flow Cytometry. **a,** Gating strategy for phenotyping. Gating was on singlets that were CD19^+^ or CD20^+^ and CD3-CD8-CD16-Ova-. Anti-IgG, IgM, IgA, IgD, CD71 and CD27 antibodies were used for B cell phenotype analysis. Antigen-specific cells were detected based on binding to RBD WT-PE^+^ and RBD WT/KEN (K417N/E484K/N501Y)-AF647^+^. **b-c,** Flow cytometry plots showing the frequency of **b,** RBD WT-binding memory B cells, and **c,** RBD-binding memory B cells cross-reactive with WT and K417N/E484K/N501Y mutant RBD in 5 selected individuals, after prime, 1.3 months, and 5 months post-second vaccination. **d,** Flow cytometry plots showing frequency of RBD-binding plasmablasts, in 10 selected vaccinees after prime or 1.3 months post-boost. **e,** Gating strategy for single-cell sorting for CD20+ memory B cells (top panel) or CD19+CD20- plasmablasts (bottom panel) which were double positive for RBD-PE and RBD-AF647. **f-g,** Representative flow cytometry plots showing dual AlexaFluor- 647-RBD and PE-RBD-binding, single-cell sorted B cells from **f,** 6 individuals after prime and 1.3 months or 5 months post-boost and **g,** 5 individuals from 1.3- or 5-months post-boost. Percentage of RBD-specific B cells is indicated.

**Extended Data Fig 3:**
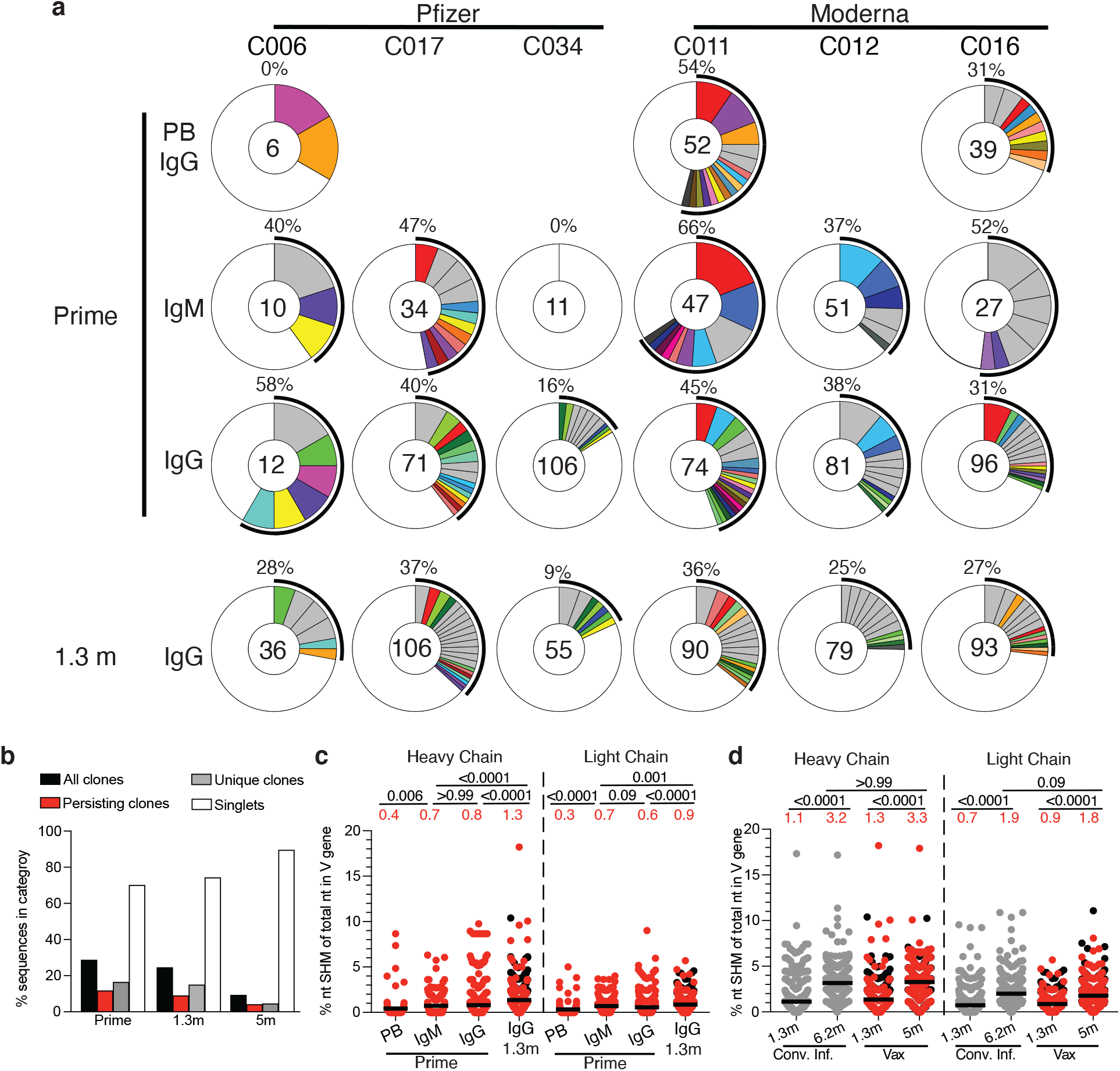
anti-SARS-CoV-2 RBD-specific plasmablast and memory B cells responses after vaccination. **a,** Pie charts show the distribution of antibody sequences from 6 individuals after prime (upper panel) or 1.3 months post-boost (lower panel). Sequences derived from IgG plasmablast (PB), IgM memory B cells (MBC), and IgG MBC compartments were analyzed after prime, while only IgG MBCs were analyzed at 1.3 months after boost, as indicated to the left of the plots. The number inside the circle indicates the number of sequences analyzed for the individual denoted above the circle. Pie slice size is proportional to the number of clonally related sequences. The black outline indicates the frequency of clonally sequences detected in each patient. Colored slices indicate persisting clones (same *IGHV* and *IGLV* genes, with highly similar CDR3s) found in multiple compartments and/or timepoints within the same patient. Grey slices indicate clones unique to the compartment. White indicates sequences isolated once. **b,** Graph shows the percentage of total paired-sequences analyzed at either prime, 1.3- or 5-months post-boost, that can be found as part of all clones (black bars), persisting clones (red bars), unique clones (grey bars), or singlets (white bar). **c-d,** Ratio of the number of somatic nucleotide mutations over the nucleotide length of the V gene in the Ig heavy and light chains, separately, in antibodies detected in **c,** different B cell compartments after prime or 1.3 months post-boost and **d,** 1.3 or 5 months post-boost compared to convalescent infected (grey) individuals after 1.3^3^ and 6.2^7^ months post-infection (also Supplementary Table 4). Horizontal bars and red numbers indicate mean ratio in each compartment at each time point. Sequences derived from samples without a prime value are shown in black. Statistical significance in **c** and **d** was determined using a Kruskal Wallis test with subsequent Dunn’s multiple comparisons.

**Extended Data Fig 4:**
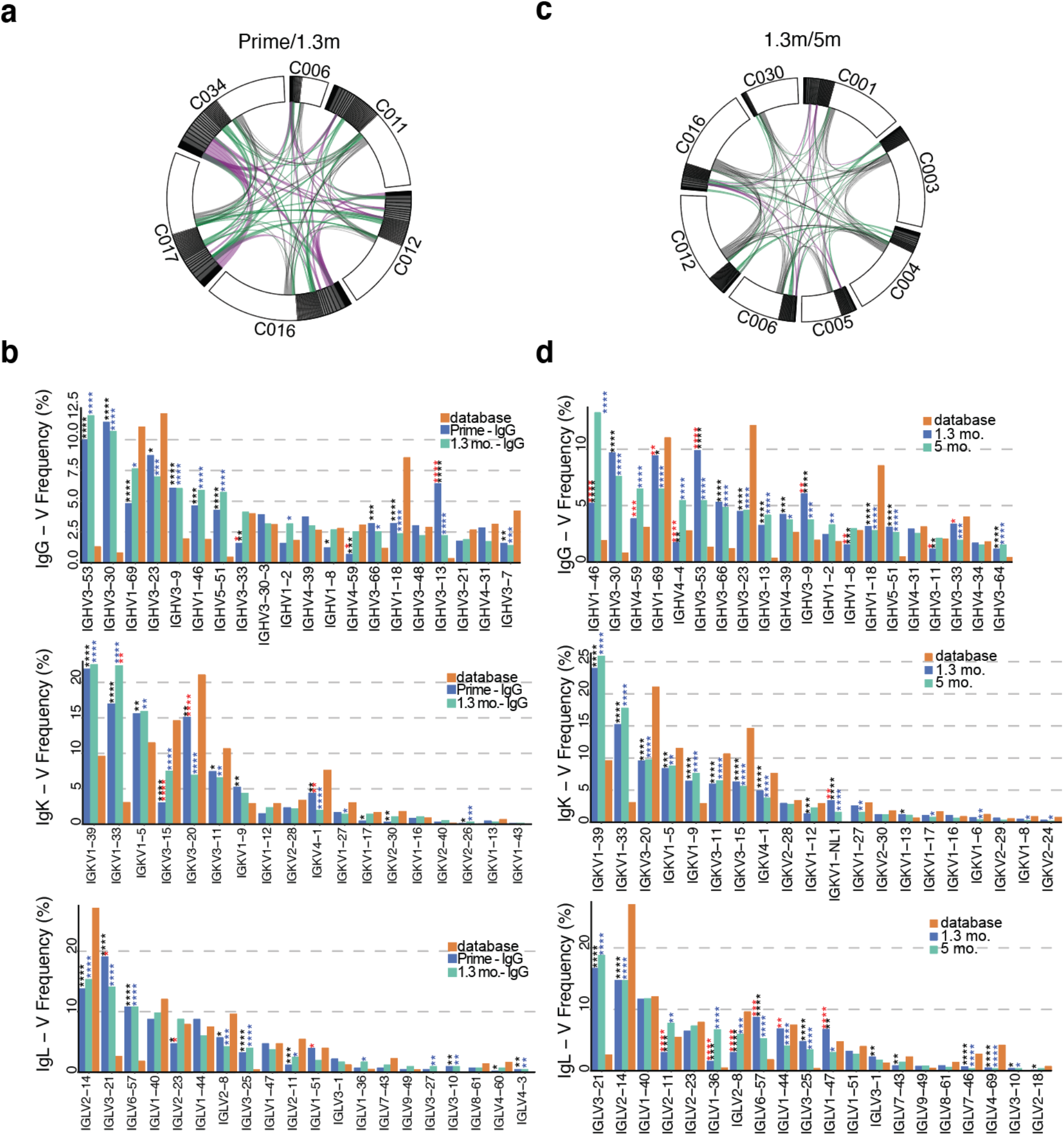
Frequency distribution of human V genes. **a,** Circos plot depicting relationship between antibodies that share V and J gene usage in both IgH and IgL when comparing prime/1.3m IgG MBC sequences. Purple, green, and grey lines connect related clones, clones and singlets, and singlets to each other, respectively. **b,** Graph shows relative abundance of human heavy chain *IGHV* (top), light chain *IGKV* (middle) or *IGLV* (bottom) genes comparing Sequence Read Archive accession SRP010970 (orange), and IgG MBCs after prime (blue) or 1.3 months post-boost (green). Statistical significance was determined by two-sided binomial test. * = p≤0.05, ** = p≤0.01, *** = p≤0.001, **** = p≤0.0001. Color of stars indicates: black - comparing Database versus Prime; blue - comparing Database versus 1.3m; red - comparing Prime versus 1.3m. **c,** Circos plot depicting relationship between antibodies that share V and J gene usage in both IgH and IgL when comparing 1.3 m/5 m IgG MBC sequences. Purple, green, and grey lines connect related clones, clones and singlets, and singlets to each other, respectively. **d,** Graph shows relative abundance of human heavy chain *IGHV* (top), light chain *IGKV* (middle) or *IGLV* (bottom) genes comparing Sequence Read Archive accession SRP010970 (orange), and IgG MBCs after 1.3 months (blue) or 5 months (green) post-vaccination. Statistical significance was determined by two-sided binomial test. * = p≤0.05, ** = p≤0.01, *** = p≤0.001, **** = p≤0.0001. Color of stars indicates: black - comparing Database versus 1.3 months; blue - comparing Database versus 5 months; red - comparing 1.3 months versus 5 months.

**Extended Data Fig 5:**
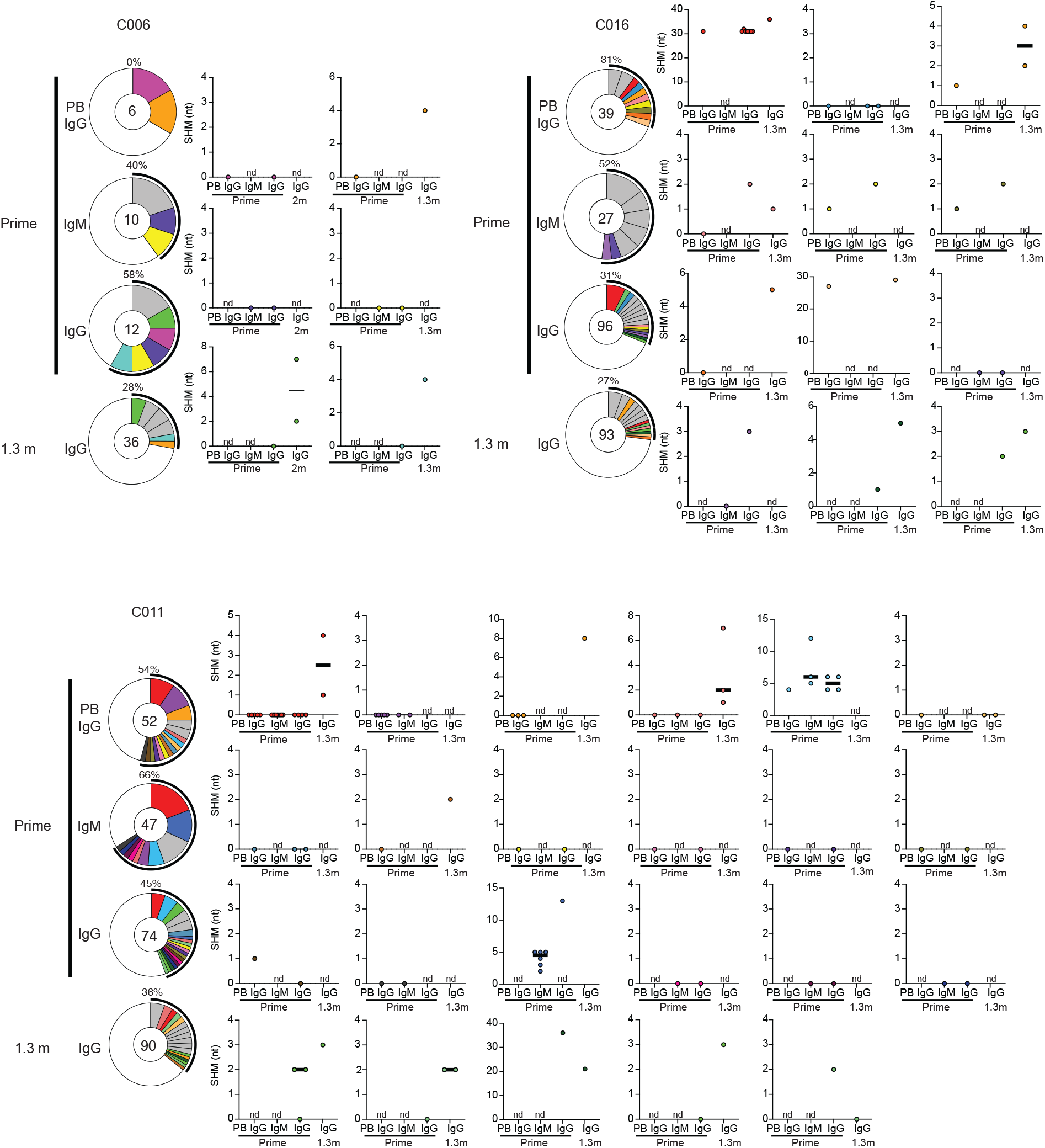
Somatic hypermutation of anti-SARS-CoV-2 RBD antibody clones after prime or boost. Clonal evolution of RBD-binding B cells from 3 individuals for which plasmablasts, IgM memory B cells, and IgG memory B cells were analyzed after prime, and IgG memory B cells were analyzed after 1.3 months post-boost (as described in Extended data Fig. 3). The number of somatic nucleotide mutations found in shared clonal families found in at least 2 different compartments is graphed to the right of each donut plot. Color of dot plots match the color of pie slices within the donut plot, which indicate persisting clones. nd – clone was Not Detected in the indicated compartment. Black horizontal line indicates median number of SHM.

**Extended Data Fig 6:**
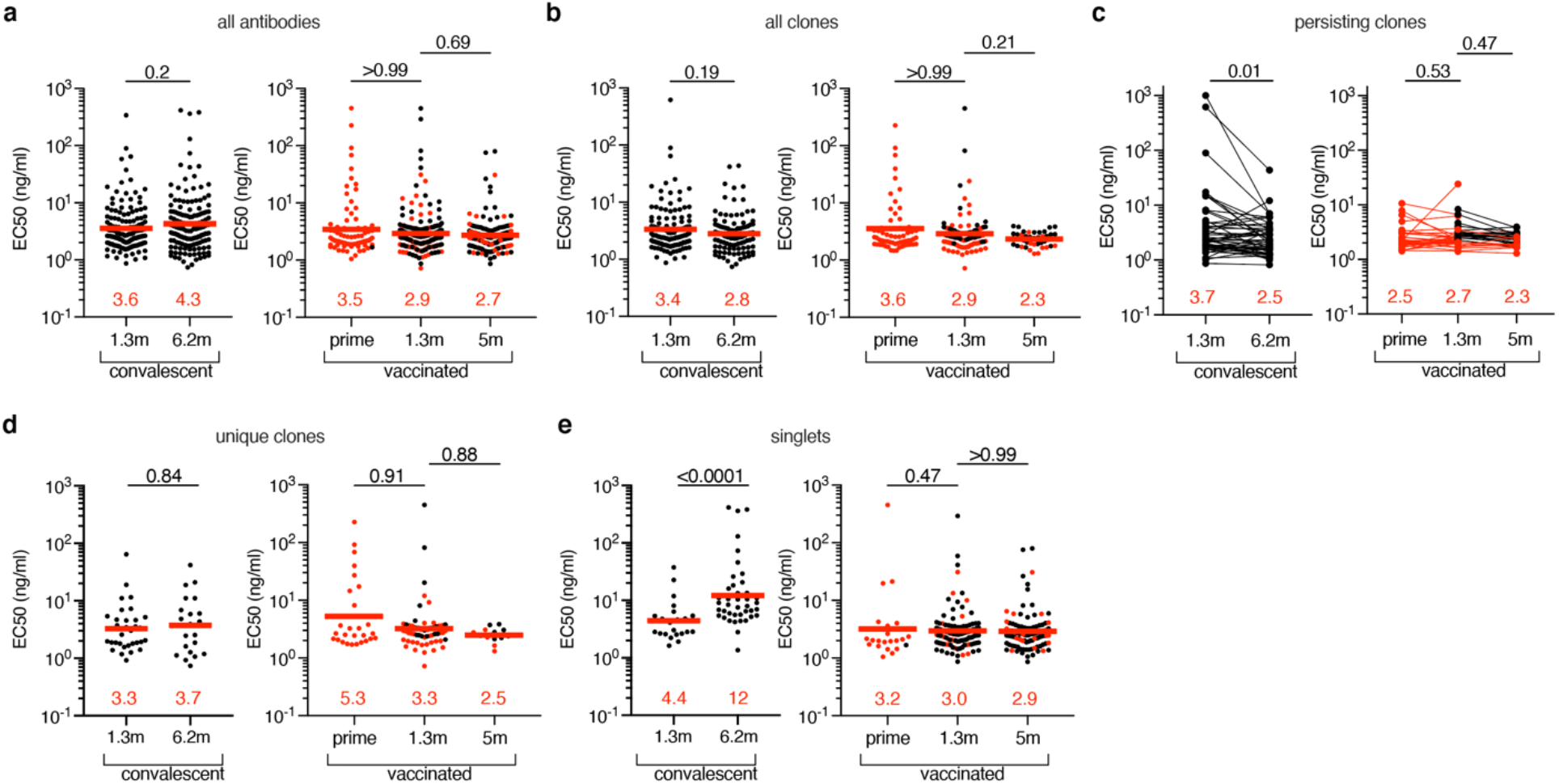
Anti-SARS-CoV-2 RBD monoclonal antibodies ELISAs. **a-e,** Graphs show anti-SARS-CoV-2 binding activity of monoclonal antibodies measured by ELISA against RBD. ELISA half-maximal concentration (EC50) values for all antibodies (**a**), all clones (**b**), persisting clones (**c**), unique clones (**d**) and singlets (**e**) isolated from COVID-19 convalescent individuals 1.3^3^ and 6.2^7^ months after infection (left panel) or from vaccinated individuals after prime, or 1.3m or 5m after receiving the second dose of mRNA vaccination (right panel). Each dot represents one antibody. Antibodies isolated from samples without a prime value are shown in black. Red horizontal bars and numbers indicate geometric mean values. Statistical significance was determined by Mann-Whitney test (left panels of **a, b, d and e**), Kruskal-Wallis test with subsequent Dunn’s multiple comparisons (right panels of **a-e**) or by Wilcoxon test (left panel of **c**). All experiments were performed at least twice.

**Extended Data Fig 7.**
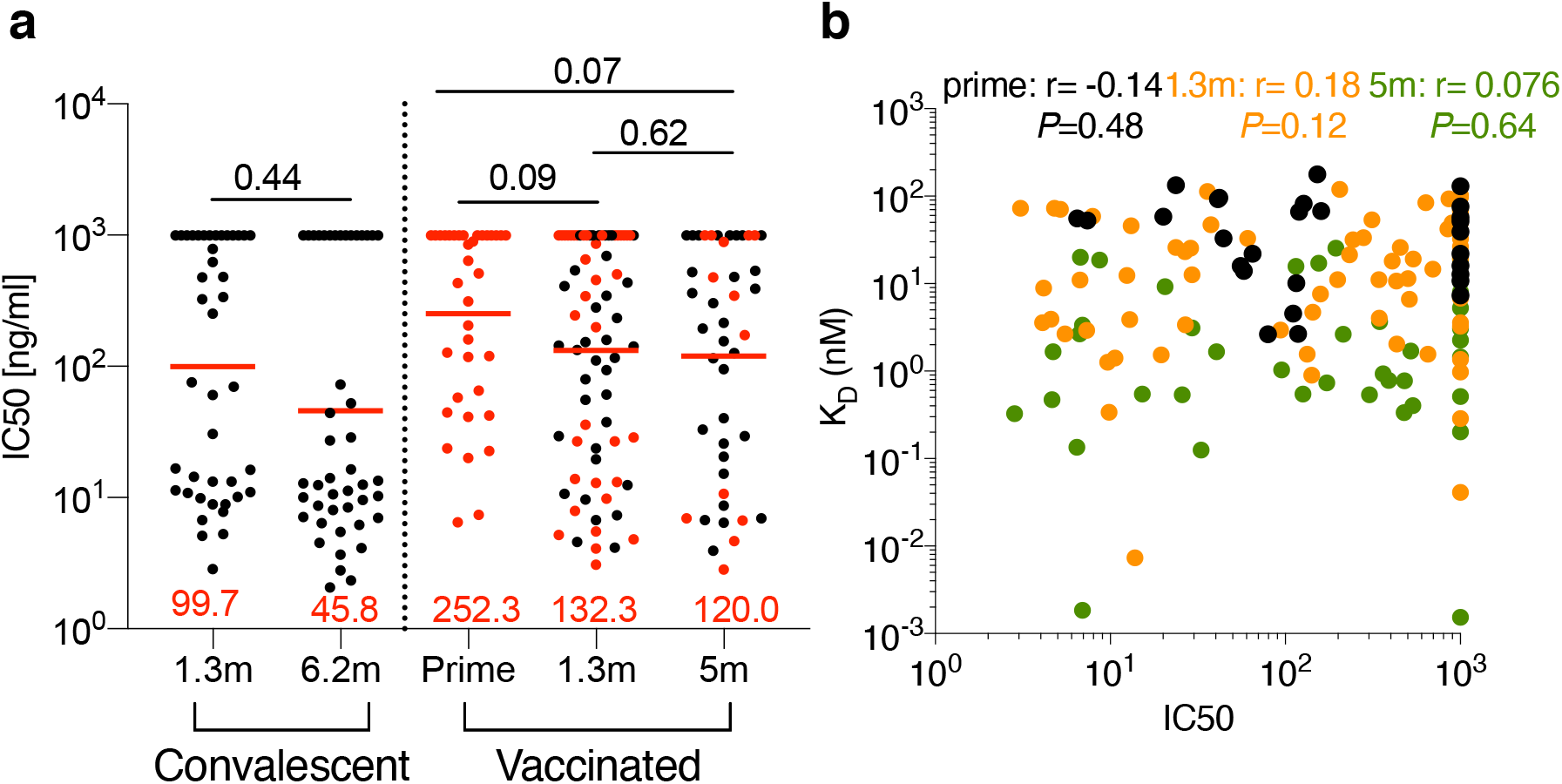
Affinity. Biolayer interferometry measurements. **a**, IC50 values for randomly selected antibodies isolated from convalescents 1.3m^3^ (n=42) and 6.2m^7^ (n=45) after infection or from vaccinees after prime (n=36), and 1.3m (n=74) and 5m (n=43). Red horizontal lines and numbers indicate geometric mean. Antibodies isolated from samples without a prime value are shown in black. **b,** Graphs show affinities (KD, Y-axis) plotted against neutralization activity (IC50, X-axis) for antibodies isolated after prime (black), or 1.3m (orange) or 5m (green) post-boost vaccination. Statistical significance was determined using Kruskal Wallis test with subsequent Dunn’s multiple comparisons in **a** and Spearman correlation test in **b**.

**Extended Data Fig 8.**
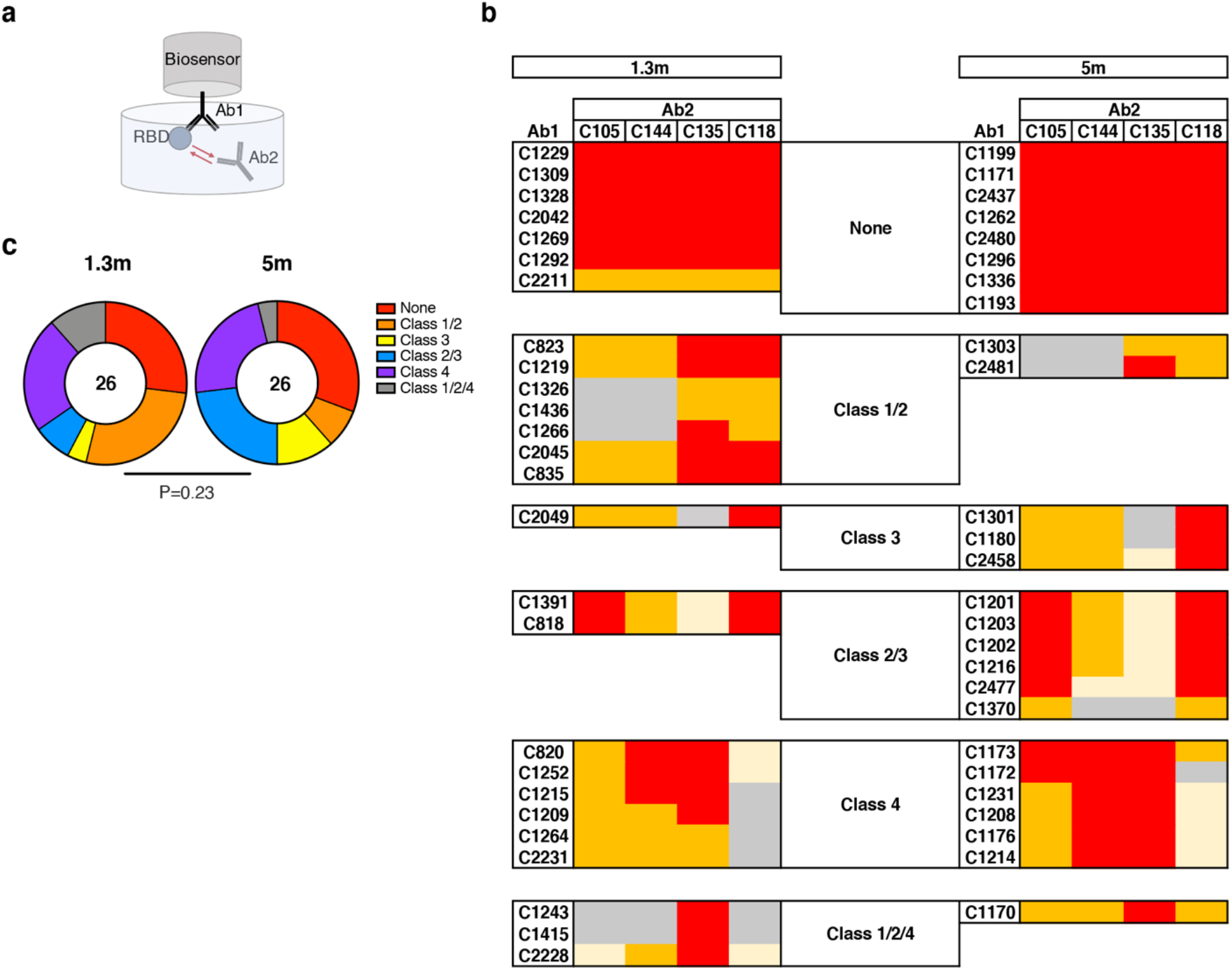
Epitope targeting of anti-SARS-CoV-2 RBD antibodies. **a**, Schematic representation of the BLI experiment for randomly selected antibodies isolated from vaccinees 1.3- and 5 months after full vaccination (each presented group shows n=26 antibodies). **b**. Heat- map of relative inhibition of Ab2 binding to the preformed Ab1-RBD complexes (grey=no binding, yellow=low binding, orange=intermediate binding, red=high binding). Values are normalized through the subtraction of the autologous antibody control. BLI traces can be found in Extended data Fig. 9. **c**. Pie charts indicate the fraction of antibodies that are assigned to different classes according to their binding pattern as shown in **b** and Extended data Fig. 9. Number in inner circle shows number of antibodies tested. Statistical significance was determined using the Chi- square test in **c**.

**Extended Data Fig 9.**
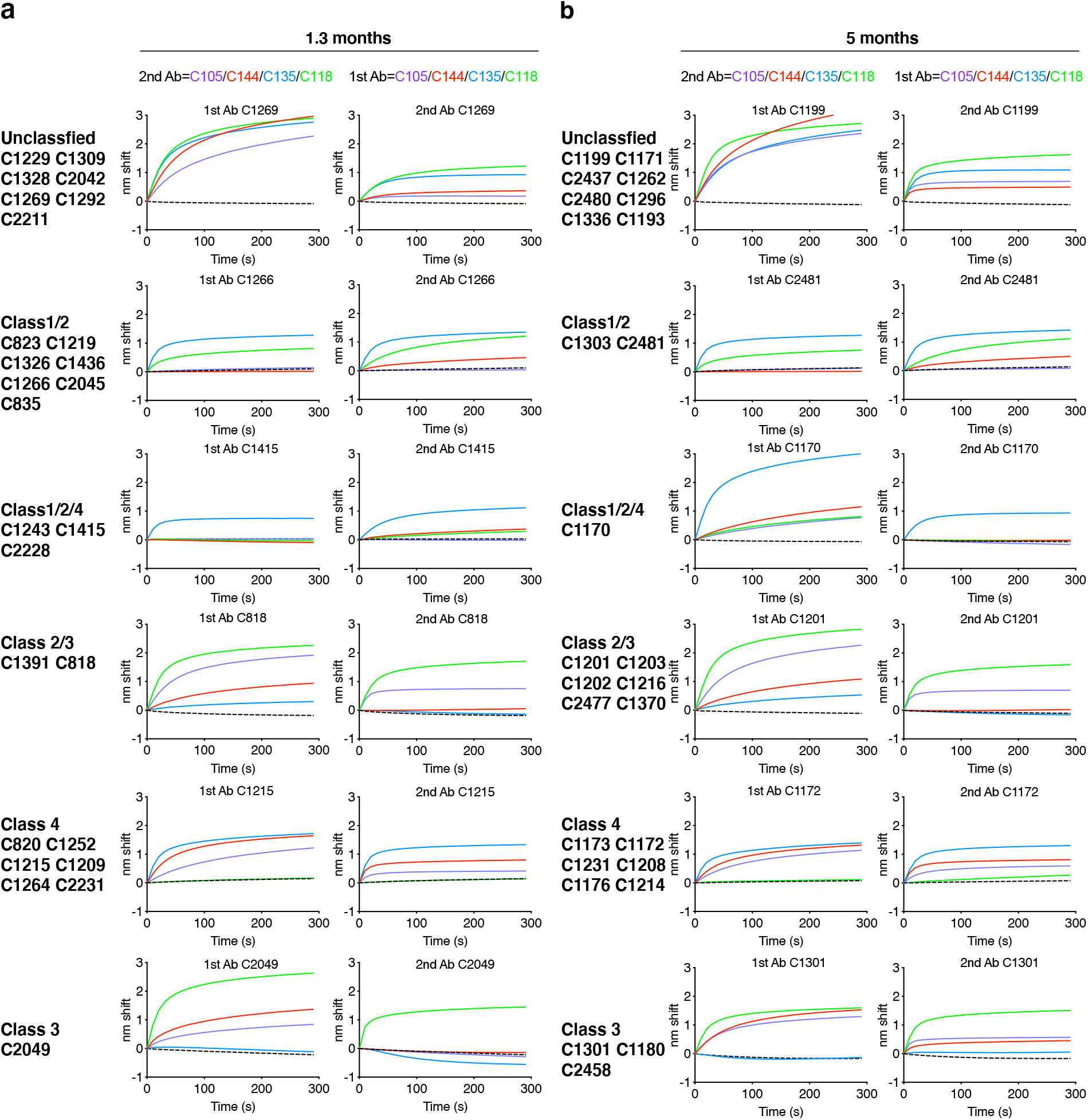
BLI traces from epitope mapping of anti-SARS-CoV-2 RBD antibodies. a-b, BLI traces from competition experiments used to determine epitope targets of anti-SARS-CoV-2 RBD antibodies isolated from vaccinees at 1.3m (**a**) or 5m (**b**) post-boost, as illustrated in Extended data Fig. 8.

**Extended Data Fig 10:**
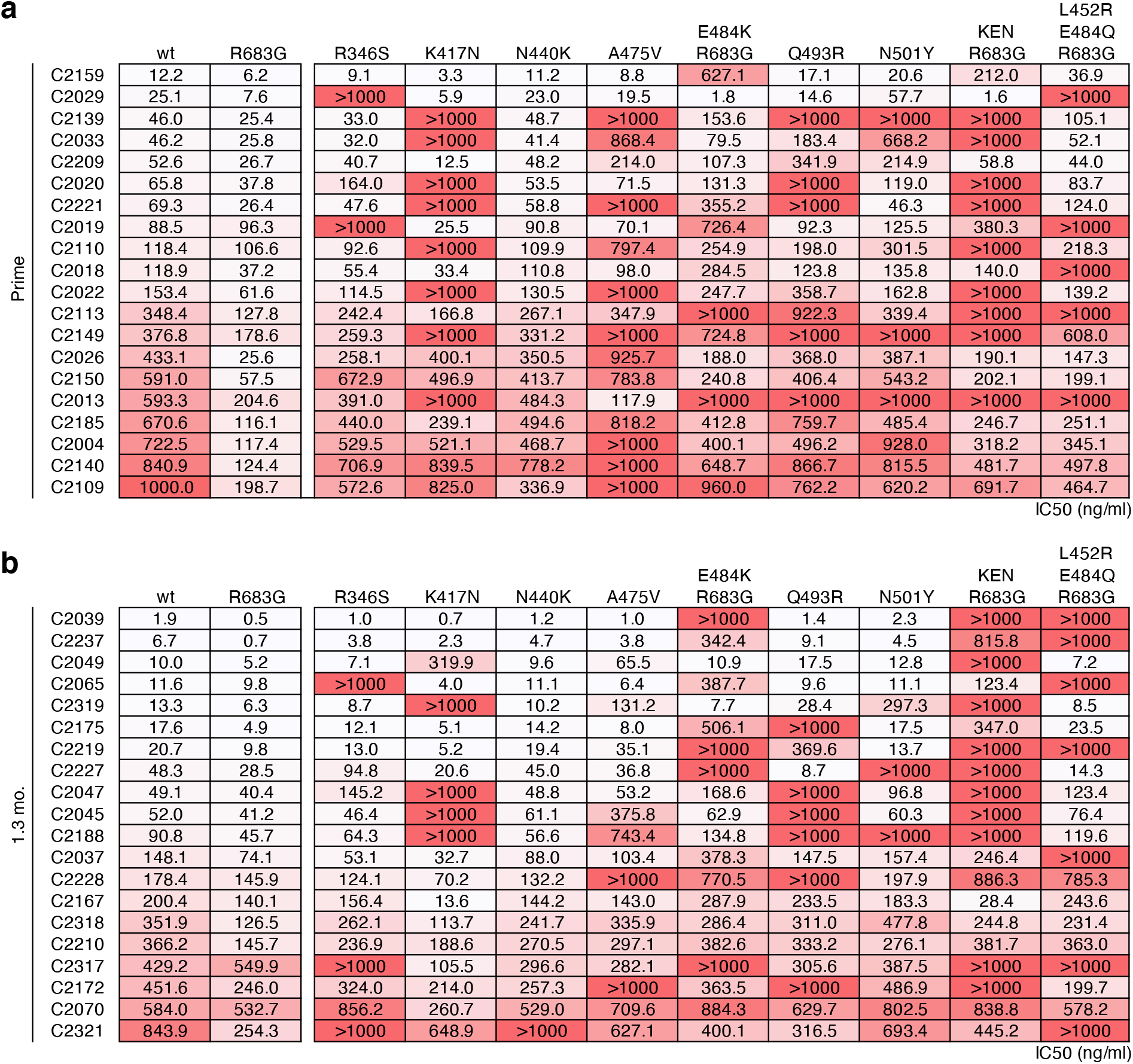
Breadth of anti-SARS-CoV-2 RBD antibodies elicited after prime and 2 doses of vaccination. **a-b**, IC50 values for n=40 neutralizing antibodies isolated after prime **(a)** or 1.3 months post-boost **(b)** against indicated mutant SARS-CoV-2 pseudoviruses. Color gradient indicates IC50 values ranging from 0 (white) to 1000 ng/ml (red).

